# Genetic variants in TMPRSS2 and Structure of SARS-CoV-2 spike glycoprotein and TMPRSS2 complex

**DOI:** 10.1101/2020.06.30.179663

**Authors:** Ravikanth Vishnubhotla, Naveen Vankadari, Vijayasarathy Ketavarapu, Ramars Amanchy, Steffie Avanthi, Govardhan Bale, Duvvur Nageshwar Reddy, Mitnala Sasikala

## Abstract

SARS-CoV-2, a highly transmittable pathogen has infected over 3.8 million people around the globe. The spike glycoprotein of SARS-CoV-2 engages host ACE2 for adhesion, TMPRSS2 for activation and entry. With the aid of whole-exome sequencing, we report a variant rs12329760 in *TMPRSS2* gene and its mutant V160M, which might impede viral entry. Furthermore, we identified TMPRSS2 cleavage sites in S2 domain of spike glycoprotein and report the structure of TMPRSS2 in complex with spike glycoprotein. We also report the structures of protease inhibitors in complex with TMPRSS2, which could hamper the interaction with spike protein. These findings advance our understanding on the role of TMPRSS2 and in the development of potential therapeutics.

## Introduction

The pandemic COVID-19 caused by a highly transmittable pathogen SARS-CoV-2, has infected millions of people worldwide^1,2^. To date, it has infected more than 3.8 million people resulting in ∼265,000 deaths with varied incidence and mortality rates across the globe^5^. The USA and European countries recorded a large number of infections associated with higher mortality compared to Asian countries. These variations could be due to differences in the virulence, pathogenicity of viral strains and host factors including genetic makeup.

SARS-CoV-2 use Spike glycoprotein for host cells adhesion. The N-terminal S1subunit interact with receptor ACE2 It is also known that SARS-CoV-2 S-protein gets activated through host proteases^7,8^. Evidences have also shown that corona viruses S proteins are cleaved by host cell proteases such as furin at S1/S2 cleavage site exposing S2 to serine protease TMPRSS2 to activate fusion of membranes for viral entry^7,12^. However, lack of structural and molecular studies concerning TMPRSS2 interaction with S-protein limits our understanding of key priming action of TMPRSS2 for the host cell entry of the S-protein, which has great therapeutic implications.

Here, with the aid of whole exome sequencing we discover and report the functionally relevant variants in *ACE2* and *TMPRSS2* in human genome that contribute to differences in COVID-19 infection rates. By employing structural mutation studies, molecular dynamics we show this variant V160M decreased stability, with a plausible role in viral entry. Furthermore, for the first time we demonstrate the structural interactions between TMPRSS2 and S2 subunit of SARS-CoV-2 spike protein. We also report the structural interactions of clinically approved protease inhibitors to block the TMPRSS2 and its further interaction with S2 subunit of spike protein establish the specificity and precision.

## Methods

### Study subjects and Ethics

The study group were recruited after obtaining informed consent and the study protocol was approved by Institutional ethics committee. Whole blood (3ml) was collected from 547 healthy individuals from the general population of Indian ethnicity. Whole exome data was generated for 20 individuals. Variants identified in *ACE2* and *TMPRSS2* were replicated in 159 and 500 individuals respectively as shown in Supplementary Fig. S6. The study subjects were considered healthy based on no self-reported disease, normal BMI, ultrasound, laboratory parameters and were found to be clinically healthy.

### DNA Isolation, Whole exome sequencing, genotyping in healthy individuals and insilico analysis

DNA was isolated from whole blood and Complete exonic regions were amplified and sequenced on the Next generation sequencer (Ion Proton; Life technologies, USA). Functionally relevant variants were identified using Polyphen and SIFT scores. Genotypes were interpreted using Genome Lab GeXP software (v10.2). All the protocols conformed to standard kit instructions and the methodology is given in Supplementary information.

### Structural modelling of TMPRSS2 and SARS-CoV-2 spike glycoprotein trimer

To better understand the structure of TMPRSS2, including the position and organization of the catalytic site for substrate processing, we modelled the monomer structure of human TMPRSS2 employing SWISS-MODEL and I-TASSER. The structure ensure all residues are placed in Ramachandran favored positions using Coot (www.mrc-imb.cam.uk/) and validated the model. we used a previously published and validated model structure of full length SARS-CoV-2 spike glycoprotein^19^.

### Molecular docking of activated form of SARS-CoV-2 glycoprotein and TMPRSS2

Molecular docking with HADDOCK 2.2 and Cluspro, interaction studies were performed with the activated form (post furin cleavage) of SARS-CoV-2 spike glycoprotein (S2 domain) and our modelled and validated TMPRSS2 (aa145-aa492) as template structures. The binding free energies were taken into consideration for selecting the best possible models. Ensured the residues occupied Ramachandran favored positions using Coot (www.mrc-imb.cam.uk/). The final model of complex structure of activated SARS-CoV-2 spike glycoprotein homotrimer and TMPRSS2 was visualized in PyMol.

### Molecular Docking and dynamics of TMPRSS2 inhibitors and structure refinement

TMPRSS2 and potent protease inhibitors (Chemostat, Upmoastat, Nafamostat and Bromhexine hydrochloride) were docked and simulated individually for molecular docking and dynamics studies. Both protein and protease inhibitors were prepared for docking by ensuring the presence of all hydrogen atoms and water molecules at least 5 Å around the binding site or catalytic pocket using the Mastero package program^34^. Binding is corroborated based on the Solvent accessibility surface area (SASA), C-Score (confidence score) and Z-score (clash score). Further jelly body refinement was done in the CCP4 program suite^35^ and then Coot (www.mrc-imb.cam.uk/) to ensure appropriate docking and no steric hindrance clashes in the side chain residues. Binding free energies were taken into consideration for selecting the best possible docked site.

## Results

### Exome Data identifies functionally relevant variants in *ACE2* and *TMPRSS2*

Whole exome sequencing on Next generation sequencer yielded 6.5-7.5 GB data and ∼56,000 variants per sample across the genome. We identified 3 variants in *ACE2* and 9 in *TMPRSS2* applying relevant filters (Fig. 1). Of these 12 variants, rs971249 in *ACE2* and rs12329760 in *TMPRSS2* genes were replicated. The minor allele frequency for the *TMPRSS2* variant was 0.22, and *ACE2* variant was 0.3.

**Figure 1:**
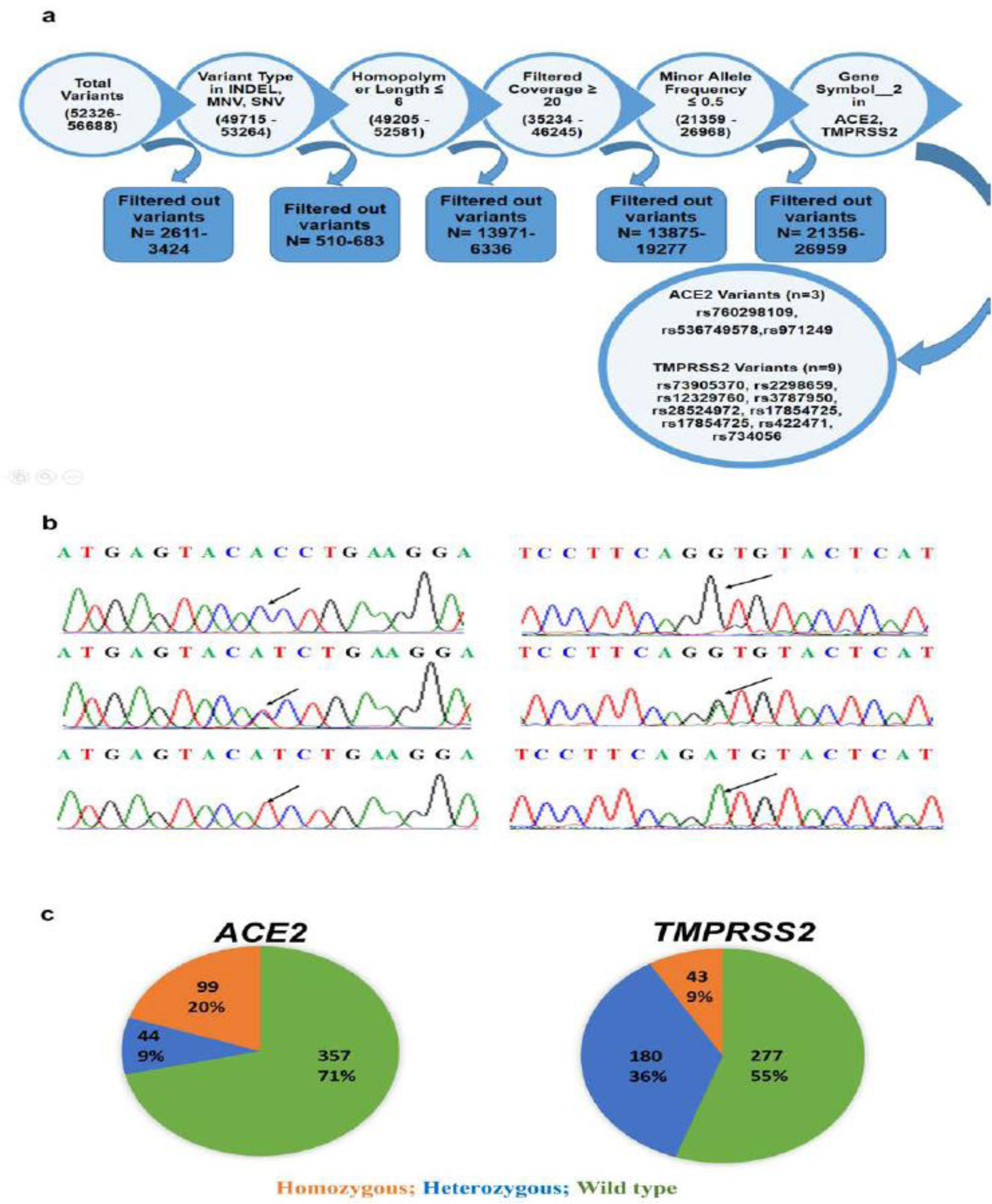
Pipelines used to identify functionally relevant variants in *ACE2* and *TMPRSS2* genes from Whole Exome sequencing data. The complete data was first filtered for quality, minor allele frequency and then for variants in both the genes. b Representative electropherogram depicting the three genotypes for the variant rs12329760 in TMPRSS2 gene.

### Allelic frequency of *ACE2* and *TMPRSS2* variants

To compare the allelic frequencies of Indians with other ethnicities we sequenced the flanking regions of the variants in *ACE2* and *TMPRSS2* in the study subjects. A representative electropherogram for *TMPRSS2* genotypes is shown (Fig. 1) The minor allele frequency (MAF) of the *ACE2* variant (rs971249-T) was 0.24 and *TMPRSS2* variant (rs12329760-T) was 0.27. the *TMPRSS2* variant was noted between ethnicities (USA, Europe Vs India; Z score= -2.08; p=0.03).

T is the minor allele. Sequence in forward strand and reverse strands with arrows indicate the variant. c. Genotype frequencies in the healthy individuals.

### Structure of TMPRSS2 and catalytic site

The overall structure of TMPRSS2 (aa145-aa492) (Fig. 2a) measures 42 Å in length and 24 Å in diameter comprising an N-terminal (aa145-aa243) activation domain and C-terminal (aa256-aa492) proteolytic or catalytic domain. We mapped the catalytic site/pocket of TMPRSS2, where residues H296, S441, K432, W461 and Q438 are found to be highly conserved with other TTPs (type II transmembrane serine proteases).

**Figure 2.**
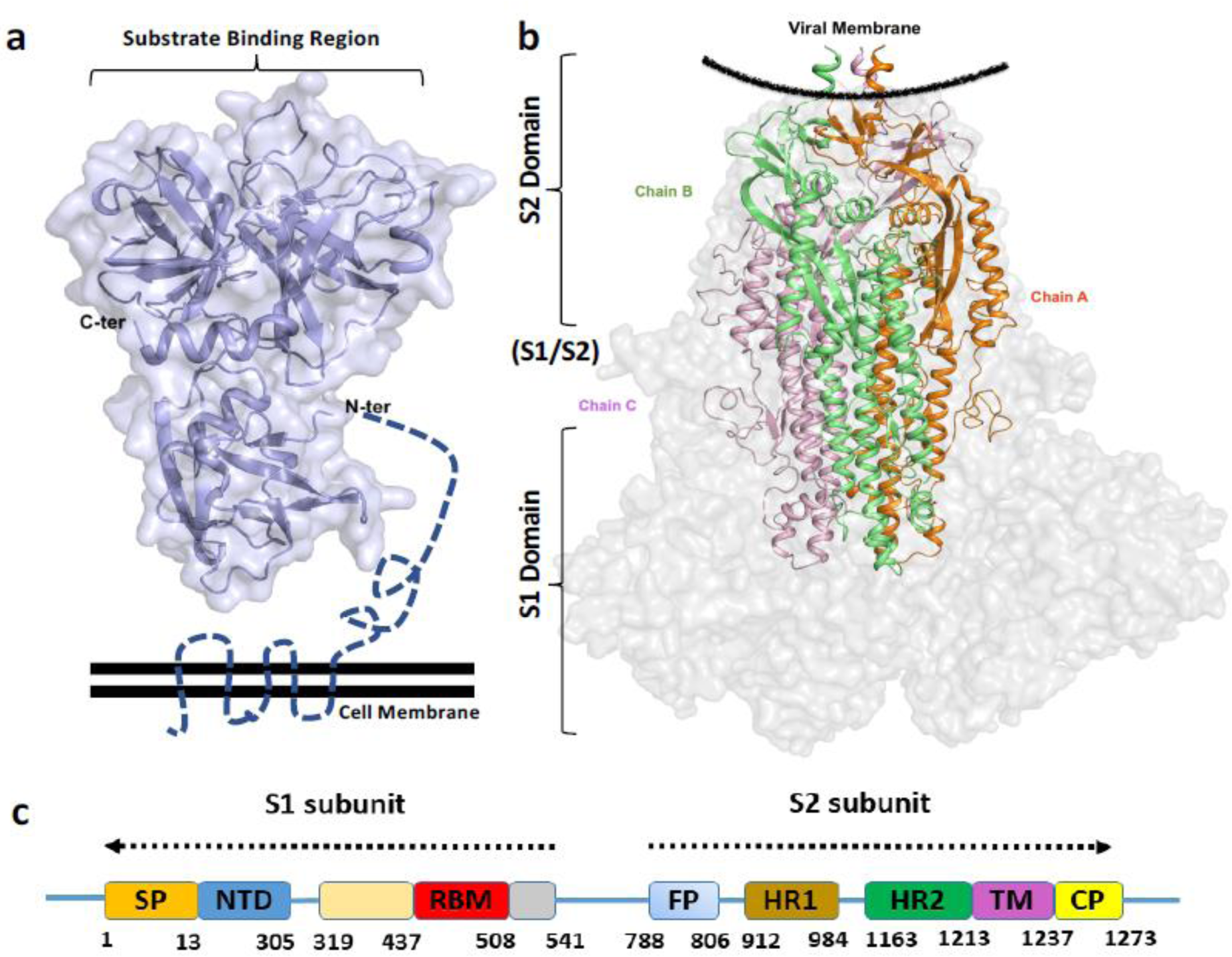
Structures of TMPRSS2 and SARS-CoV-2 spike glycoprotein (a) Surface and ribbon model showing the side view of homology model structure of TMPRSS2. The substrate or catalytic binding region on the apical region is labelled and the unstructured and membrane region is shown in dashed lines. (b) Surface and ribbon model showing the side view of SARS-CoV2 spike glycoprotein. The S1 domain region is shown is grey surface and the internal S2 domain is shown coloured ribbon. The three protomers of homotrimer are colored accordingly.(c) Domain arrangement of of S1 and S2 subunits of Spike glycoprotein SP: Signal peptide; NTD: N-terminal domain; RBM: Receptor binding domain; FP: Fusion peptide; HR1: Heptad repeat1; HR2: Heptad repeat2; TM: Transmembrane domain and CP: Cytoplasmic domain.

### Identification of TMPRSS2 binding sites in SARS-CoV-2 spike glycoprotein and structure of activated S-protein

Based on available literature addressing the serine protease target sites we mapped two potential TMPRSS2 binding sites in S-protein (T1: aa837 to aa 845 and T2: aa 976 to aa986) located in the S2 domain region (Fig. 3). The structure of S-proteins with S1 and S2 domains are shown in (Fig. 3b).

**Figure. 3.**
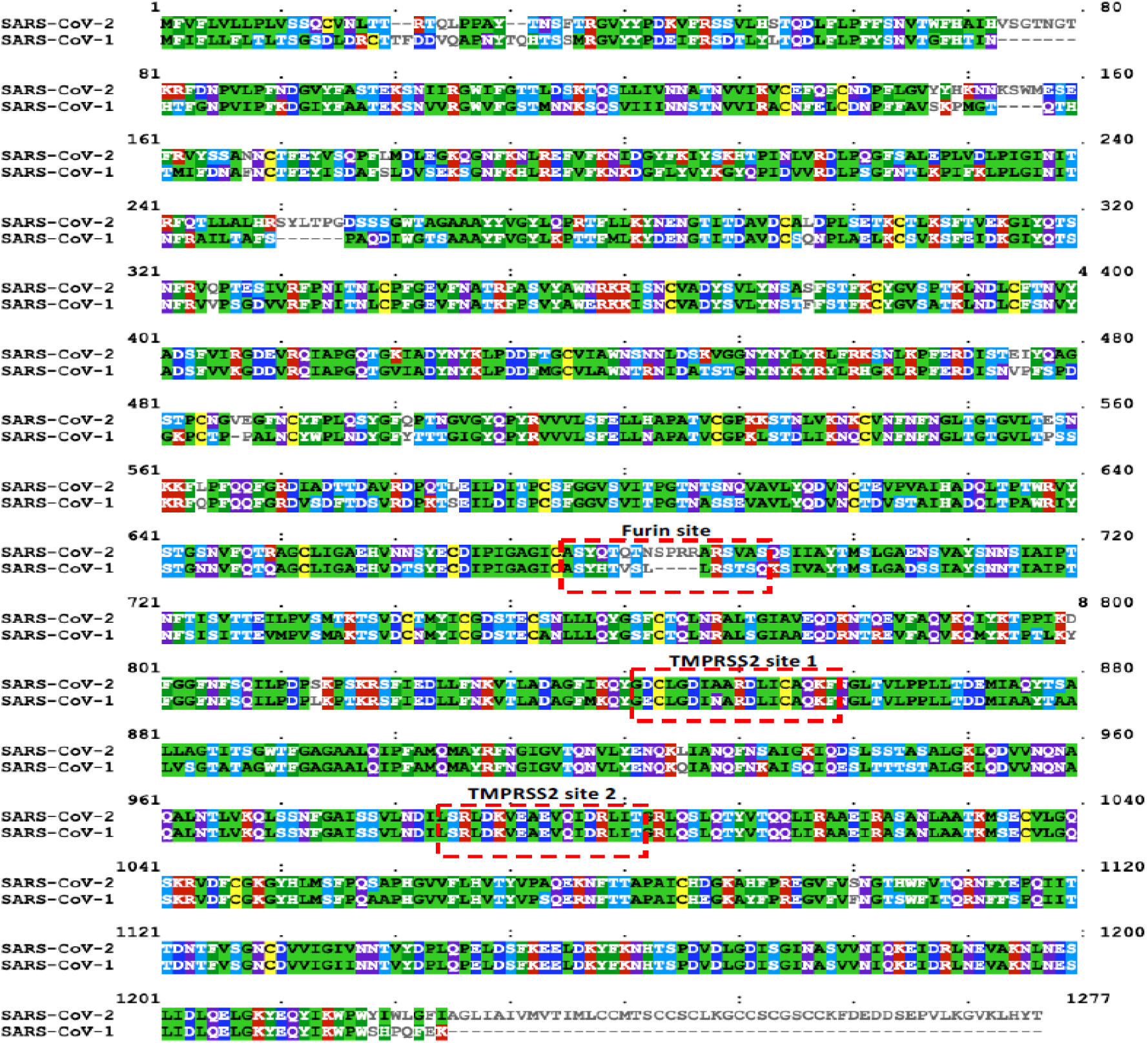
A ClustalW amino acid sequence alignment showing identical, conserved and non-conserved (unique) residues (NOT shaded) of full length SARS-CoV-2 and SARS-CoV-1 Spike glycoproteins. The respective Furin and TMPRSS binding sites are labelled and marked accordingly.

### V160M decreases the stability of TMPRSS2

The TMPRSS2 variant (rs12329760) and the current epidemiological reports showing lower SARS-CoV-2 infection rate in the Indian ethnic background raises an intriguing question about its correlation with the infection. Valine 160 in the wild-type protein (V160) is located in the N-terminal SRCS domain (scavenger receptor cysteine rich) and is paired with the three anti-parallel β-sheets (aa145-aa170), Structurally the replacement of V160M does not accommodate the Met due to the topology and charge limit (Fig. 4a,). We performed molecular dynamics and simulation studies to know mutant protein characteristics and found undergone structural deformation due to the complete shift in the motif (Fig.4c,). This is corroborated with the overall increase in the B-factor (stability factor) of TMPRSS2 with V160M mutation. (Fig. 4 e, f). These observations reiterate that the V160M variant of TMPRSS2 decreases the stability of the protein.

**Figure 4.**
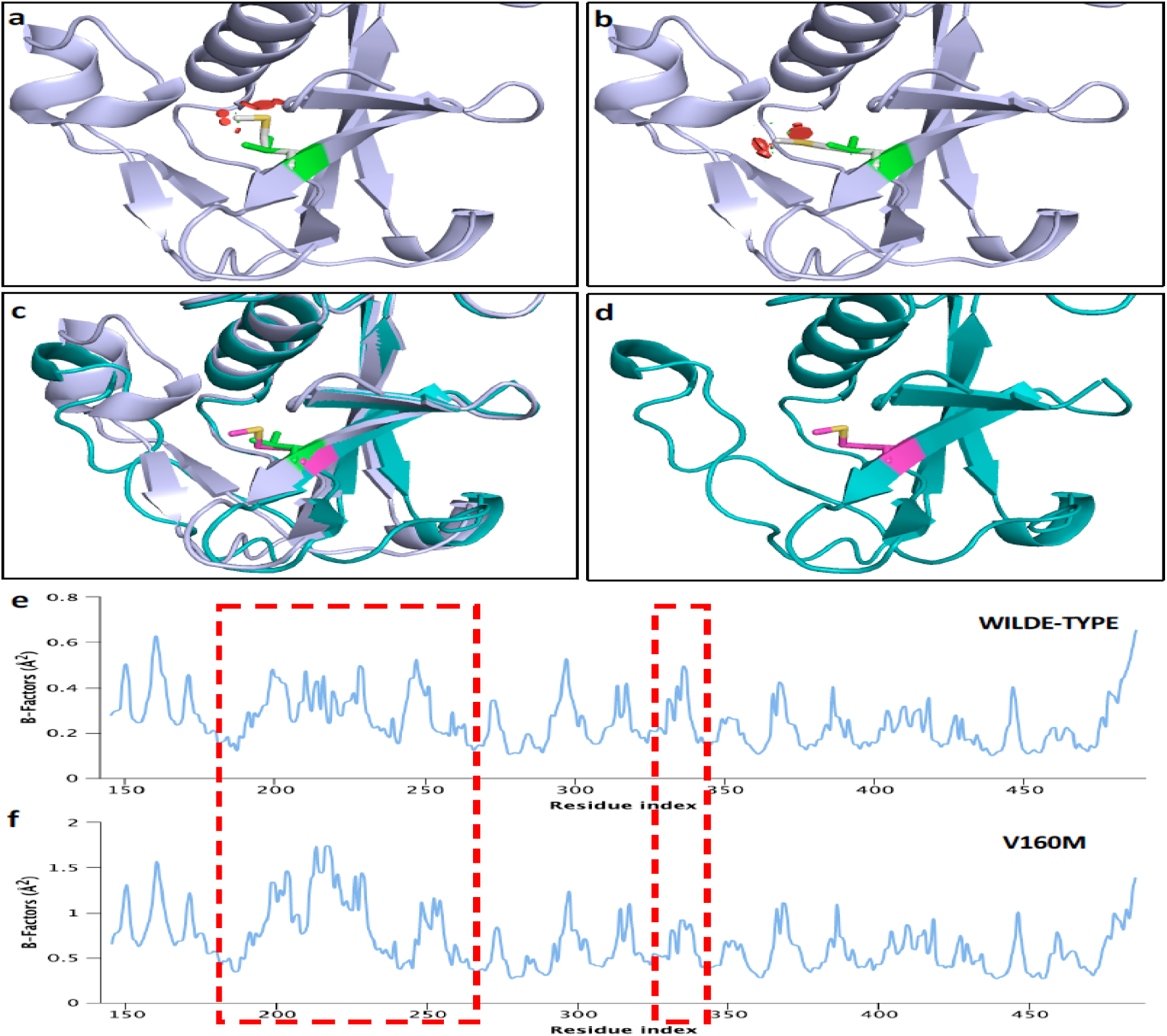
TMPRSS2 genetic variant V160M and its structural changes (a and b) Cartoon representation of N-terminal domain TMPRSS2 (blue) structure. The native V160 is shown in green stick and replacement with M160 shown in the same location resulting clash (red dots) is noticeable. (c) Superimposition of native (blue) and V160M mutant (slate) structure of TMPRSS2. (d) 3D structure V160M mutant TMPRSS2. The native V160 is shown in green and mutant M160 is shown in pink sticks. (e and f) B-factor profile of wildtype and V160M mutant structure of TMPRSS2. The difference in the domain oscillation and structural change is marked in red dashed box.

### Structural interactions in the activated SARS-CoV-2 glycoprotein - TMPRSS2 complex

We have observed TMPRSS2 potentially interacting with the T1 and T2 sites of the S2 domain of spike glycoprotein. The overall complex structure shows that TMPRSS2 binds to T1 site of SARS-CoV-2 spike glycoprotein homo-trimer and adopts typical protease binding mode and interacts with a higher affinity (−250 kcal). This suggests a bonafide and tight interaction of TMPRSS2 protease with the spike glycoprotein. The active or catalytic pocket of TMPRSS2 is made of residues Q276, H296, E299, K300, P301, K340, K342, E389, K390, L419, S441, Q438 and W461. And the residues H296, S441, K432 and Q438 are highly conserved among several serine proteases. The TMPRSS2 catalytic pocket of a cup-like architecture accommodates and interacts with the linker region connecting α1/α2 and α3/α4 of S2 domain of the spike protein Fig. 5a, 5b. With respect to TMPRSS2 interaction with T2 site of activated S2 domain of SARS-CoV-2 spike glycoprotein, the extended and well-exposed loop region (G832 to N856) adopts the peptide binding mode with the TMPRSS2. The entire “S-shape” loop region of T2 site (K835 to I850) of S2 domain pass into the canyon-like crevice or cup-like structure of TMPRSS2 catalytic pocket (Fig. 5d) Among them, residues K835, Y837, D839, C840, L841, D843, I844, R847, R848 are the key interacting residues and also well positioned towards the catalytic pocket of the TMPRSS2 (Fig. 5 e).

**Figure 5.**
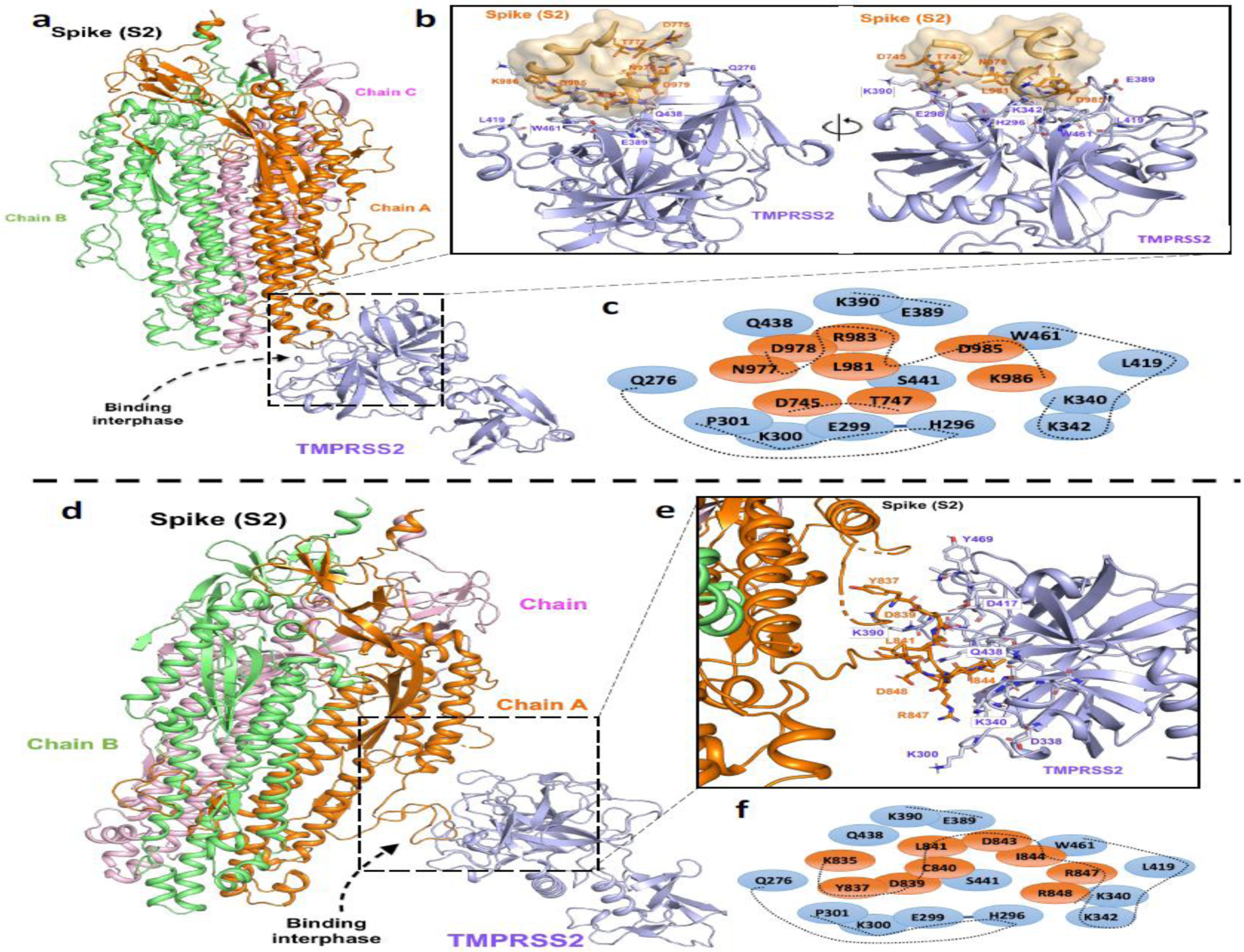
Two binding modes of interaction between TMPRSS2 and S2 domain of spike glycoprotein (a) T1: First model of TMPRSS interacting with S2 domain of spike glycoprotein at T1 region (b) Enlarged view showing the detailed interaction between TMRPRSS2 and T1 site and S2 domain. (c). Key residues and binding orientation topology at the interaction interface at T1 site. (d) T2: Alternate model of TMPRSS interaction with S2 domain of spike glycoprotein at T2 region. (e) Enlarged (front and orthogonal) view showing the detailed interaction between TMRPRSS2 and T2 site and S2 domain (f) Key residues and binding orientation topology at the interaction interface at T2 site. Colour coding and labelling was retained same in all figures

### Structure of TMPRSS2 in complex with clinically approved protease inhibitors

To explore the specificity and validate the precision of the catalytic pocket we performed molecular docking, refinement and dynamic studies with four potential TMPRSS2 protease inhibitors (Chemostat, Upamostat, Nafmostat and Bromhexine hydrochloride) (Fig. 6 a-f). As expected we observed the specific binding location of all drugs directed to the catalytic pocket of TMPRSS2. The main drug binding region is structurally linked with the binding of T1 and T2 sites of the S2 domain of spike glycol proteins with binding energy of -9 kCal.

**Figure 6.**
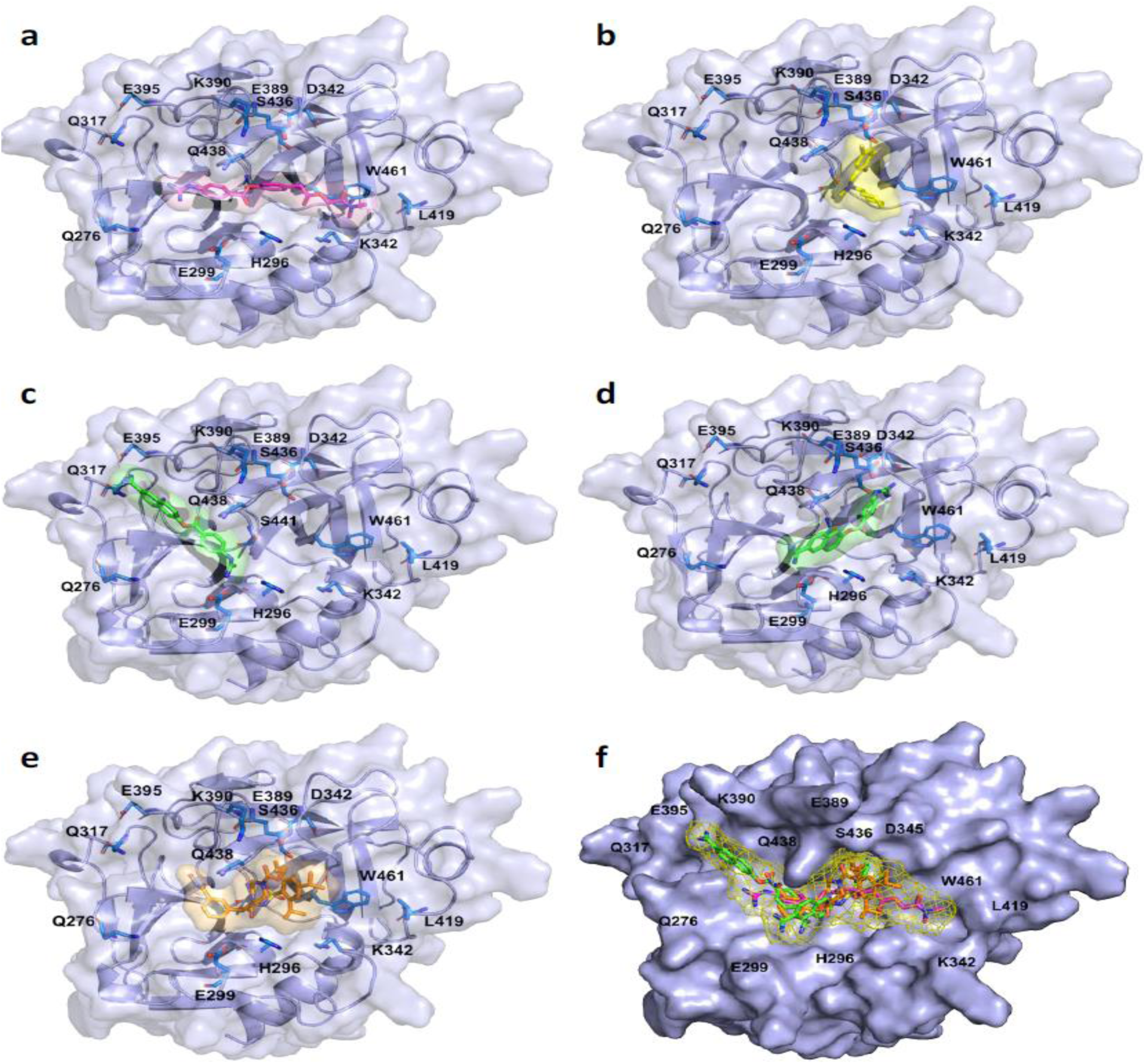
Surface and cartoon representation structures of TMPRSS2 with bound potential protease inhibitors. The TMPRSS2 structure is shown in blue and the active site residues are labelled accordingly. The bound inhibitor (a) Camostat (b) Bromhexine (c) and (d) Nafamostat binding in two different modes (e) Upamostat (f) Superimposition of all four inhibitors in the catalytic pocket of TMPRSS2

We next analyzed interactions of individual drugs with TMPRSS2 catalytic pocket of active residues Q276, H296, E299, K300, P301, K340, K342, E389, K390, L419, S441, Q438 and W461. It was interesting to notice that all four inhibitors (Chemostat, Upamostat, Nafamostat and Bromhexine hydrochloride) predominantly bind to the specific location of the TMPRSS2 catalytic pocket. The detailed position and alignment of amino acids of TMPRSS2 and individual drugs involved in the interaction are showed in Fig. 7.

**Figure. 7.**
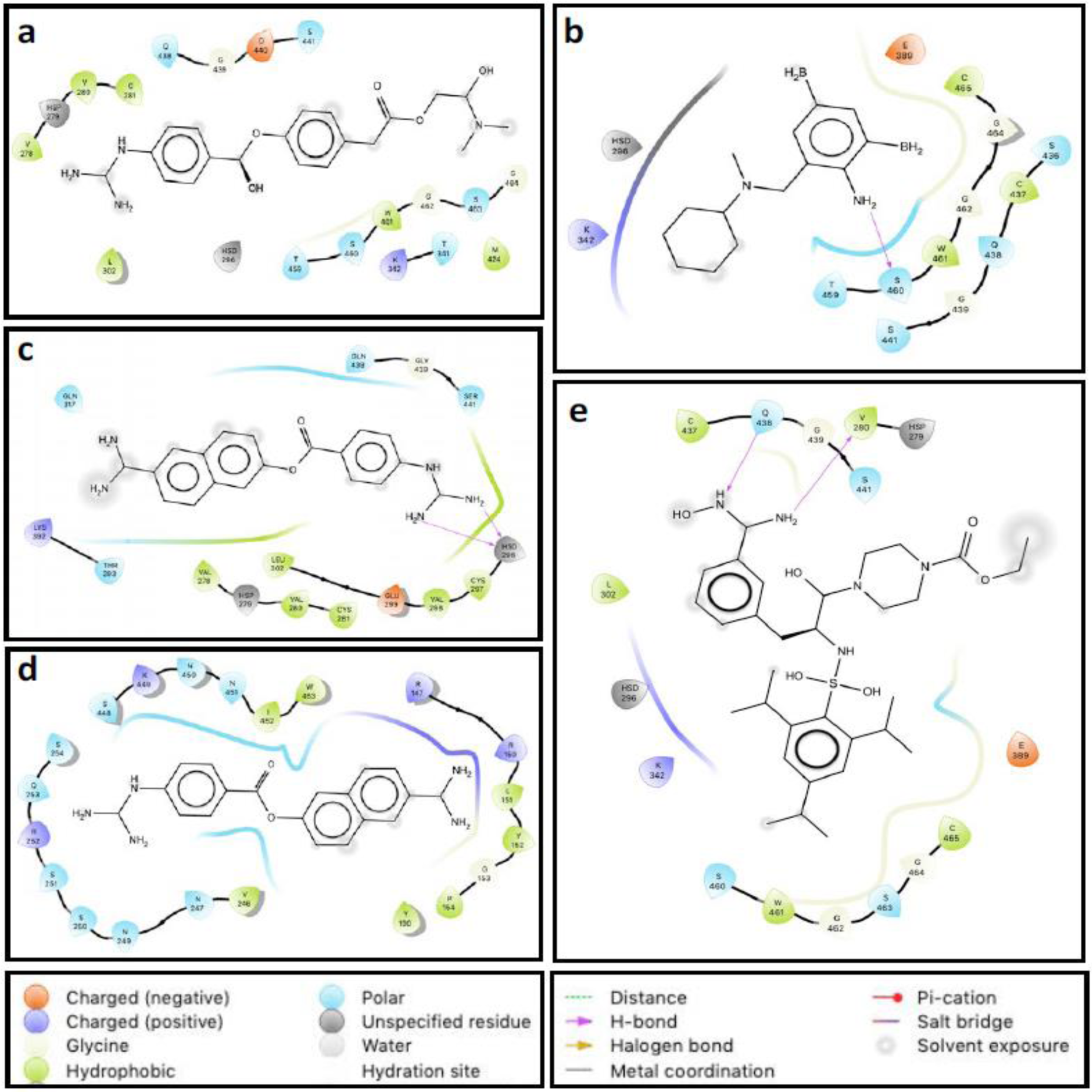
Detailed structural view of the interaction between different potential protease inhibitors and TMPRSS2 catalytic site. The position and residue names are labelled accordingly and type of interaction between the individual drug and surrounding amino acids are marked as shown in the figure key. (a) Camostat (b) Bromhexine (c) and (d) Nafamostat binding in two different modes (e) Upamostat

## Discussion

We identified a missense variant rs12329760 in *TMPRSS2* gene that is shown to confer decreased stability of the protein. From the homology model of TMPRSS2 and its mutant V160M, we have identified its interaction sites with SARS-CoV-2 S-protein for the first time. Binding of clinically proven protease inhibitors with precision and specificity at these interaction sites suggest that this site can be targeted for developing newer drugs in the treatment of SARS-CoV-2 infection and till then these drugs can be tested on patients therapeutically in current COVID clinics.

It is intriguing to note that, although the viral strain emerged from China and spread to various regions of the world, significant differences in the incidence and mortality rates cannot be completely explained by the various quasi-sub species of the virus. This suggests that the host factors *ACE2* receptor and *TMPRSS2*, play a role in the infection. Significant differences (p=0.03) in allelic frequency of *TMPRSS2* variant across ethnicities (MAF and variant carriers 0.27; 44.60% in Indians, 0.22; 36.26% in Europeans, 0.15;28.24% in Americans), indicate its role in varied infectivity. The identified variant rs12329760; V160M in *TMPRSS2* gene was earlier reported to have a higher risk of TMPRSS2-ERG fusion in prostate cancer patients^18^. It also gains importance as it is known to be present in an exonic splicing enhancer site (Srp40) that is associated with an increased chance of exon skipping or malformation of the protein due to potential exonic splicing enhancer site disruption ^13,14^. Hence, we believe that the presence of unstable protein in the variant carriers might hamper the entry of the virus in to the host.

Despite knowing the role of TMPRSS2 in priming of S-protein for host cell entry, its structure and interaction sites with the S-protein are not elucidated. Also the fact that a deleterious variant in *TMPRSS2* gene is identified in healthy individuals, we wanted to discern its position in relation to the activation site employing a validated homology model. To understand the structure/functional relevance of the variant in *TMPRSS2*, the precise location of the variant on the protein was explored. This predicted and validated model was important because the structure for TMPRSS2 protease was not available in PDB and none of the previous studies demonstrated the protease cleavage sites.The homology model clearly demonstrates that the variant V160M is in the N-terminal domain (aa145-aa243) towards cell membrane away from catalytic and C-terminal domain (aa256-aa492) located outwards.

We have also identified the interactions of TMPRSS2 with spike glycoprotein using the above model in complex with the validated model structure of full-length SARS-CoV-2 spike glycoprotein^19^ as the available structures with PDB lack Furin and TMPRSS2 recognition and cleavage sites ^11^. Based on available literature on the target sites of other serine proteases ^20,21^ along with sequence analysis, we identified two prime TMPRSS2 recognition sites in the SARS-CoV-2 spike glycoprotein. The two identified sites span a part of heptad region1 (HR1) of six helix bundle of SARS-CoV-2 near the fusion peptide. Recently it is reported that heptad regions HR1 and HR2 aid in bringing the fusion peptide in close proximity to transmembrane domain facilitating membrane fusion^22-24^. It is convincing to speculate that Furin protease acts first on the spike glycoprotein at S1/S2 region and cleaves the spike protein to S1 (ACE2 and CD26 binding region) and S2 (trimerization domain) resulting in the complete exposure of S2 domain and TMPRSS2 recognition sites as elucidated earlier ^12^,^25^.

While this model fits well for the normal genotype of TMPRSS2, whether genetic variant in TMPRSS2 protease has any physiological role in SARS-CoV-2 infection or activation of spike glycoprotein is elusive. Insilico studies suggested decreased stability of the protein in variant, as the free energy change is less than -0.5 k cal/mol, and is present in an important domain of the protein ^15,26^. Molecular dynamics simulation confirm structural distortions due to complete shift in the motif leading to changes in quaternary structure of the protein and concomitant increase in B factor with V160M variant. This reiterates the fact that although the V160M is not located in the catalytic pocket of TMPRSS2, decrease in the stability of the protein might hamper SARS-CoV-2 viral entry, however, this needs to be confirmed.

In order to assess the binding efficiency of TMPRSS2 with identified cleavage sites on S2 subunit of Spike protein, we tested the clinically proven protease inhibitors: Camostat, Nafamostat, Upamostat and Bromhexine hydrochloride (BHH). Camostat used therapeutically, for unrelated clinical conditions was shown to inhibit influenza viral replication^27,28^ while Nafamostat is a potent inhibitor of MERS S-protein mediated membrane fusion ^29^. BHH on the other hand is FDA approved mucolytic agent and a specific inhibitor of TMPRSS2 ^30,31^. Upamostat is another serine protease inhibitor under consideration for clinical trials ^32^. All four drugs bind to the active site of TMPRSS2 with high precision and specificity (Fig. 5 and Supplementary Fig. S5). We demonstrate that Camostat, Upamostat and BHH preferentially bind to a specific location at the catalytic pocket of TMPRSS2, while Nafamostat binds to 3 additional binding residues in two modes.Furthermore, these potential TMPRSS2 inhibitors also share their interaction via several polar, charged and hydrophobic interactions (Supplementary Fig. S5). The specific binding of these drugs with high precision confirms that they could potentially bind and impede the interaction between the spike glycoprotein and TMPRSS2. This study identified the interaction sites of the drugs with TMPRSS2 and therefore, this conserved epitope can be targeted for developing vaccines and therapeutic drugs. Based on these findings we propose a model of interaction of TMPRSS2 with S2 subunit of spike protein of SARS-CoV-2 Fig. 8. Limitation of this study is that we have not shown the association of V160M variant with infectivity in COVID-19 patients and instability of the protein is shown by *in silico* approaches. Further studies are needed in COVID-19 patients to explore the association and functional assays to gain mechanical insights on the role of the variant.

**Figure 8.**
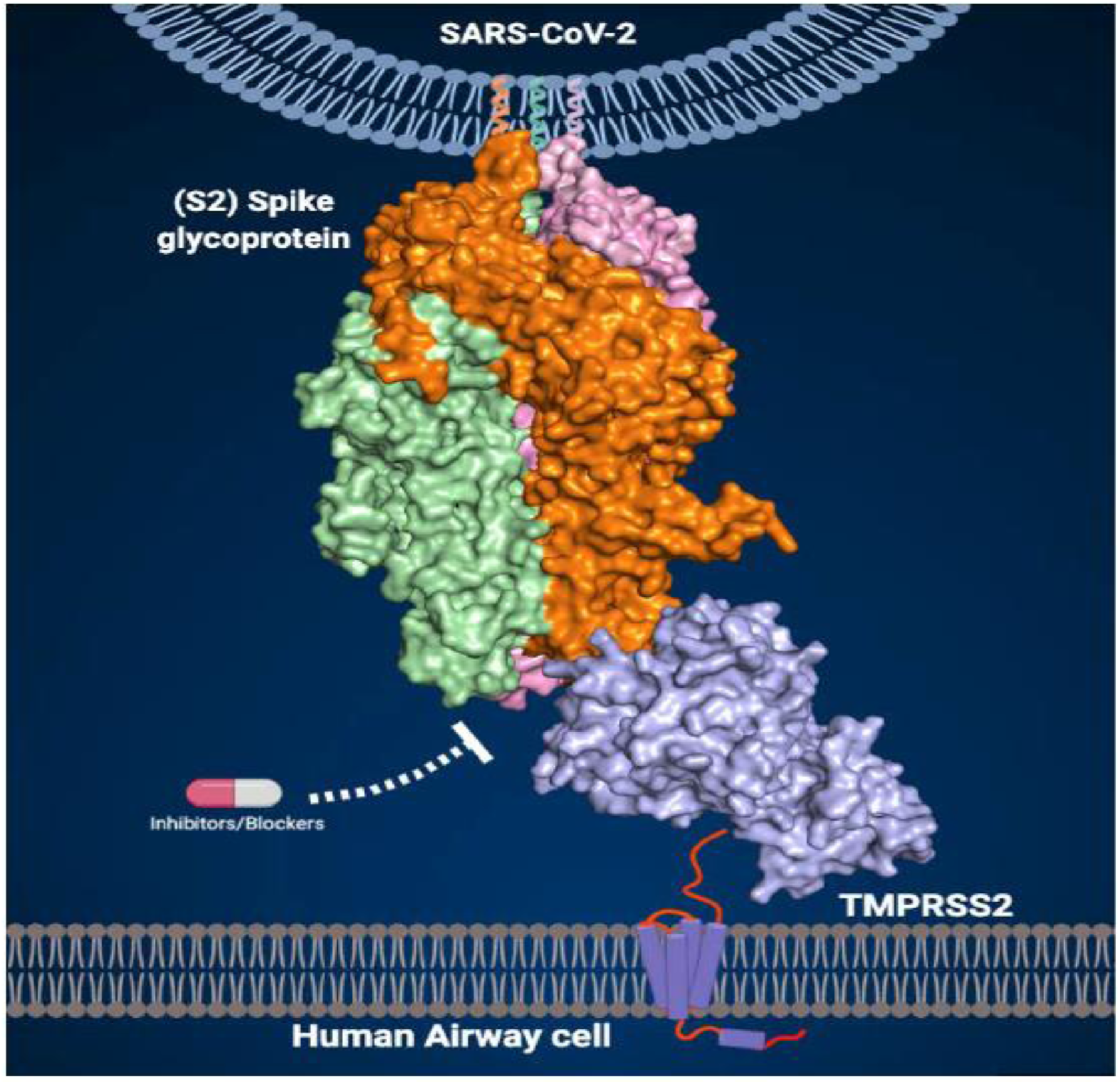
Proposed Model of TMPRSS2 binding S2 unit of SARS-CoV-2 spike glycoprotein. The three protomers of spike protein are coloured accordingly.

As in other human diseases, the incidence, severity, progression and therapeutic response have a genetic predisposition, susceptibility to COVID-19 may also have a predisposition. Although age, sex and comorbidities are shown to associate with varied incidence and severity of the disease, host genetic predisposition has not been explored. This study reports for the first time host factor genetics, molecular structure of TMPRSS2 and its cleavage sites on S2 subunit of spike glycoprotein. It is interesting to note that a significant proportion are variant carriers in India with lesser incidence of COVID-19 as compared to other ethnicities including USA and European countries that have a lower variant carriers with higher incidence. While we report the importance of host genetic factors for the first time, COVID-19 host genetics initiative (https://www.covid19hg.org/) reiterate the need to study the role of human genome in explaining COVID-19 severity and susceptibility.

## Acknowledgements

The authors acknowledge the funding received from Asian Healthcare Foundation. We thank the Monash University Software Platform for license and access to the concerned softwares. RA would like to thank the Science and Engineering Research Board (SERB), Government of India for grant support (EMR/2016/005994).

## Author Contributions

RV performed analysis and interpretation of whole exome sequencing, genotyping data, insilico analysis and drafted the manuscript; NV performed Homology modelling of TMPRSS2, docking studies for TMPRSS2 – S-protein and drug interaction in TMPRSS2, Molecular dynamics for structural interactions, simulation of TMPRSS2 variant, interpretation of data and drafting the manuscript; VK performed Homology modelling of TMPRSS2 and its variants, screening of protease inhibitors from literature, drug bank and drafted the manuscript; RA contributed to proofreading and revising the manuscript; SA and GB extracted the DNA, performed whole exome sequencing and genotyping; DNR Conception of the study and directed the research with critical inputs and revision of the manuscript; MS conception of the study, designing and supervising the experiments, participated in discussions with NV at Monash University, interpreted the data, drafted and revised the manuscript

